# Recurrent Vulvovaginal Candidiasis; a dynamic interkingdom biofilm disease of *Candida* and *Lactobacillus*

**DOI:** 10.1101/2021.02.12.430906

**Authors:** Emily McKloud, Leighann Sherry, Ryan Kean, Christopher Delaney, Shanice Williams, Rebecca Metcalfe, Rachael Thomas, Craig Williams, Gordon Ramage

## Abstract

Vulvovaginal Candidiasis (VVC) is the most prevalent *Candida* infection in humans affecting 75% of women at least once throughout their lifetime. In its debilitating recurrent form, RVVC is estimated to affect 140 million women annually. Despite this strikingly high prevalence, treatment options for RVVC remain limited with many women experiencing failed clinical treatment with frontline azoles. Further, the cause of onset and recurrence of disease is largely unknown with few studies identifying potential mechanisms of failed treatment. This study aimed to assess a panel of clinical samples from healthy women and those with RVVC to investigate the influence of *Candida*, vaginal microbiome and antagonism between *Candida* and *Lactobacillus* on disease pathology. 16S rRNA sequencing characterised disease by a reduction in specific health-associated *Lactobacillus* such as *L. crispatus*, coupled with an increase in *L. iners*. *In vitro* analysis showed *Candida albicans* clinical isolates are capable of heterogeneous biofilm formation and show the presence of hyphae and *C. albicans* aggregates in vaginal lavage. Additionally, the ability of *Lactobacillus* to inhibit *C. albicans* biofilm formation and biofilm-related gene expression was demonstrated. Using RNA sequencing technology, we were able to exploit a possible mechanism by which *L. crispatus* may aim to re-establish a healthy vaginal environment through amino-acid acquisition from *C. albicans*. This study suggests RVVC is not entirely due to an arbitrary switch in *C. albicans* from commensal to pathogen and understanding interactions between the yeast and vaginal *Lactobacillus* species may be more crucial to elucidating the cause of RVVC and developing appropriate therapies.

## Importance

Fungal infections are becoming increasingly recognised as a substantial health burden on the global population. Over 1 billion people are estimated to suffer from fungal infections each year, resulting in over 1.5 million deaths [1]. These infections are commonly associated with mucosal sites such as the vagina, gut and oral cavity. Vaginitis is estimated to account for up to 7% of all visits to gynaecologists and for up to 10 million general practitioner (GP) appointments annually [2]. VVC is not a reportable disease and is often self-treated using over-the-counter antifungal agents, subsequently the exact prevalence and distribution is impossible to determine. VVC is reported as the second most common cause of vaginitis and it is estimated that 75% of women will suffer from VVC in their childbearing years with up to 140 million of these women developing recurrent VVC (RVVC) annually, defined as ≥4 cases within one year [3, 4]. These recurrent cases are debilitating, impact quality of life and can result in significant health implications including pelvic inflammatory disease, infertility and menstrual disorders. VVC infections have been linked to use of antibiotics and contraceptives as well as new sexual partners, however it is not considered a sexually transmitted disease and no distinguishable cause has been identified [5, 6].

The biofilm-forming yeast *Candida albicans* is reported as the predominant pathogen responsible for up to 90% of VVC infections [4]. *C. albicans* is commonly isolated from the vagina as an asymptomatic commensal with a carriage rate of up to 33% in pre-menopausal women [7]. Other non*-albicans Candida* (NAC) species, including C. glabrata, C. parapsilosis, C. dublinensis and C. krusei account for 10-20% of VVC infections and are associated with complicated VVC, with more severe symptoms and higher recurrence rates [8]. Isolated species in NAC VVC infections include *C. glabrata, C. krusei, C. parapsilosis, C. tropicalis* and *C. dubliniensis,* with *C. glabrata* being the most common [9]. *Candida* species, predominantly *C. albicans*, form thick biofilms which dramatically increase fungal tolerance to drugs commonly used in the treatment of VVC such as fluconazole, miconazole and flucytosine [10]. Sessile cells have been shown to tolerate antifungal concentrations 1000-fold greater than their planktonic counterparts [11]. *C. albicans* has been shown to form biofilms on vaginal mucosa both *ex vivo* and *in vivo* by identification of significant fungal load, biofilm architecture and extra-cellular matrix using confocal and electron microscopy [12]. This is a possible contributing factor to failed clinical treatment resulting in persistent and recurrent VVC. Despite the identification of *Candida* biofilms on vaginal mucosa, the predicted therapeutic challenge this presents in VVC is still disputed. Other studies challenge the hypothesis of vaginal biofilms and instead suggest other factors such as germ tube formation and polymicrobial tissue invasion as a more critical feature of VVC [13, 14].

The vaginal microbiome is somewhat unique in that a ‘healthy’ microbiome is associated with a less diverse community of microbes, predominantly lactobacilli, namely *L. crispatus*, *L. iners*, *L. gasseri* and *L. jensenii* [15]. Dysbiosis of the vaginal microbiome leads to an increase in species diversity, with fewer lactobacilli and greater numbers of pathogenic anaerobes such as *Gardnerella* and *Prevotella* [16, 17]. In healthy women, lactic acid bacteria (LAB) are thought to be responsible for maintaining a homeostatic microbiome by inhibiting growth and adhesion of other microbes via the production of secreted metabolites such as lactic acid, biosurfactants, bacteriocins and H_2_O_2_. *L. crispatus* is a prevalent commensal of the healthy vaginal environment in various microbiome studies and produces both lactic acid and H_2_O_2_ [18, 19]. Additionally, *L. crispatus* secretes the L-lactic acid isomer which has been extensively studied for its ability to lower vaginal pH, elicit anti-inflammatory responses and inhibit microbial colonisation [20, 21]. Conversely, lactic acid lowers the pH of the vaginal environment which could result in *Candida* stress responses leading to hyphae formation and release of virulence factors. This was shown by Beyer and colleagues where the MAP kinase CgHog1 of *C. glabrata* was upregulated in response to clinically relevant concentrations of lactic acid [22]. CgHog1 increases the ability of *C. glabrata* to persist within the body, suggesting this could contribute to severity of RVVC.

A single dose of oral fluconazole is sufficient to treat sporadic *C. albicans* VVC in 80-90% of cases [23]. Treatment of VVC caused by a NAC is more complicated, requiring prolonged suppressive azole therapies and is often unsuccessful. The efficacy of therapeutics such as amphotericin B and boric acid has been assessed for the treatment of RVVC. Both drugs when delivered intravaginally for 14-21 days were found to be effective in around 70% of patients [24, 25]. An alternative suggestion is that RVVC may be managed through microbiome replacement therapy, possibly through the use of *Lactobacillus* probiotics [26]. Probiotic therapy involves the administration of live microorganisms which directly results in a health benefit for the patient [27]. Due to the diversity of lactobacilli within the vaginal microbiome, it is difficult to estimate which species would be most important to replace with probiotic therapy. One study evaluated *L. plantarum* P17630 combined with the standard treatment of clotrimazole for 3 days and concluded a potential resolution of vaginal discomfort [28].

RVVC is a significant burden both economically and for women’s health, with its prevalence is poorly documented globally due to the levels of self-treatment. Identifying triggers for development and recurrence of VVC and the pathogenesis of the microbes involved, could considerably improve prevention and treatment options for women with recurrent, azole-resistant cases. This study therefore aimed to screen a panel of clinical samples from healthy women and those with RVVC for the presence of *Candida* and to investigate microbial diversity between the two cohorts. Additionally, we aimed to utilise RNA sequencing technology to elucidate the intimate interactions of *C. albicans* and *L. crispatus* in a biofilm co-culture experiment.

## Materials and Methods

### Patient recruitment and collection of clinical samples

One hundredwomen aged 18 and over attending Glasgow Sandyford Sexual Health Clinic were enrolled in the study. Patients presenting with symptomatic RVVC (n=40) were recruited as well as asymptomatic women attending the clinic for contraceptive intrauterine device (IUD) implantation, acting as a healthy cohort (n=60). This study was granted ethical approval by the Sheffield Research Ethics Committee (16/YH/0310). Patients were excluded from the study if they had active bacterial vaginosis, were pregnant, immunosuppressed, menstruating, menopausal or had taken antibiotics/antifungals within 7 days prior to sampling. From each patient, one high vaginal swab (HVS) and one cervico-vaginal lavage (CVL) were collected. A graphical representation of sample collection and processing can be found in Supplementary Figure 1.

### Detection of inflammatory biomarkers in CVL

CVL supernatants were recovered by centrifugation for assessment of IL-8 levels by Enzyme-Linked Immunosorbent Assay (ELISA) (Invitrogen, Paisley, UK), following manufacturer’s instructions. CVL was diluted 1:2 and absorbances were measured using a spectrophotometer at 450nm and 570nm.

### Quantification of microbial load by quantitative PCR

HVS samples were used to extract DNA for quantitative PCR (qPCR) analysis of *Candida*/bacterial burden. DNA was extracted using the QIAmp DNA mini kit, as per manufacturer’s instructions (Qiagen, Crawley, UK) and qPCR used to quantify fungal and bacterial load in each sample. Primers specific to the conserved *Candida* ITS rRNA gene were used to determine *Candida* load. For bacterial load, primers and probe specific to the 16S rRNA gene were used [29]. Primer sequences can be found in Supplementary Table 1. Total qPCR reaction volume was 20 μL, with 1 μL of extracted DNA, 500 μM of forward/reverse primers and UV-treated RNase free H_2_O. For 16S, 250 μM of probe and 2X Taqman™ Universal PCR Master Mix (ThermoScientific, Loughborough, UK) was used, 2X Fast SYBR^®^ Green PCR Master Mix (ThermoScientific, Loughborough, UK) was used for ITS primers. qPCR was carried out using a Step-One plus real time PCR machine (Life Technologies, Paisley, UK) with the following thermal conditions: An activation step of 50°C for 2 min and 95°C for 10 min followed by 40 cycles of 95°C for 10 s to denature and annealing at 60°C for 30 s. Standard curves constructed from serially diluted DNA of *C. albicans* SC5314 and *Escherichia coli* K12 were used extrapolate *Candida* and bacterial colony forming equivalents (CFE/mL), respectively, as described previously [30].

### Preparation of 16S amplicon libraries for Illumina sequencing

To observe microbial populations in the samples, 16S rRNA sequencing was performed. Briefly, previously extracted HVS DNA was used to sequence the 16S rRNA V4 region using the Illumina MiSeq sequencing platform (Edinburgh Genomics) using 2×250bp paired-end reads. Amplification of the V4 region was achieved using fusion Golay adaptors barcoded on the reverse strand as described previously [31].

### Identification of Candida species from cervico-vaginal lavage

Lavage samples were screened for presence and identification of *Candida* species using Colorex *Candida* chromogenic agar (E&O Laboratories Ltd, Bonnybridge, UK) and Matrix Assisted Laser Desorption/Ionization – Time Of Flight (MALDI-TOF) mass spectrometry. For culture identification, 20 μL of CVL samples were spread across the surface of a chromogenic agar plate before 48 h incubation at 30°C. The colour of the colonies cultured was used to determine *Candida* species, colony numbers were used to calculate the number of colony forming units per millilitre (CFU/mL). Clinical isolates were then stored on beads in glycerol in Microbank^®^ vials (Pro-Lab Diagnostics, Cheshire, UK) at −80°C. Isolate identities were confirmed by MALDI-TOF analysis using a Bruker Microflex, comparing recorded spectra to the Bruker database.

### Screening Candida isolates for biofilm formation

*Candida* clinical isolates were assessed for biofilm forming capabilities by crystal violet assay. *Candida* isolates (n=33) were cultured on Sabouraud’s (SAB) agar for 48 h at 30°C. For biofilm formation, overnight cultures were grown in yeast extract peptone dextrose (YPD) at 30°C. Cultures were washed twice with PBS and standardised in RPMI-1640 medium to a final cell density of 1×10^6^ CFU/mL. Eight biofilms of each isolate were grown in 96-well, flat bottomed polystyrene microtiter plates for 24 h at 37°C before biomass measured by the crystal violet assay [32]. For visualisation of *Candida* aggregates, 30 μL of CVL was stained with calcofluor white (CFW) to a final concentration of 0.06 μg/mL for 1 h at 37°C, before being imaged using the EVOS live cell imaging system (ThermoScientific, Loughborough, UK).

### Media preparation

Todd Hewitt broth (THB, Merck UK), used in biofilm co-culture experiments, was supplemented with 10 μM menadione and 10 μg/mL hemin (ThermoFisher) and mixed 1:1 with RPMI (Herein referred to as 1:1 broth).

### Antagonism of C. albicans and Lactobacillus in co-culture

The ability of the following 7 *Lactobacillus* strains to inhibit *C. albicans* SC5314 biofilm formation was assessed: *L. casei* ATCC 393, *L. fermentum* ATCC 14931, *L. crispatus* ATCC 33820, *L. iners* DSMZ 13335, *L. salivarius* ATCC 11741, *L. jensenii* ATCC 25258 and *L. rhamnosus* ATCC 7469. For biofilm formation, overnight cultures were standardised to 1×10^6^ CFU/mL for *C. albicans* and 1×10^7^ CFU/mL for *Lactobacillus* species in 1:1 medium, as described previously. Eight biofilms of each *C. albicans*-*Lactobacillus* pair were incubated in 5% CO_2_ for 24 h. In addition, *C. albicans* biofilms were grown for 4 h prior to *Lactobacillus* being added for 20 h, with biomass quantified using the crystal violet assay. To assess *C. albicans* biofilm-related gene expression, RNA was extracted from dual species biofilms using the PureLink RNA mini kit (ThermoScientific, Loughborough, UK), following manufacturer’s instructions. In brief, 2 μg of RNA was converted to cDNA using the high capacity cDNA reverse transcription kit (ThermoScientific, Loughborough, UK) and 1 μL used in a 20 μL qPCR reaction with 10 μL 2X Fast SYBR^®^ Green PCR Master Mix, 1 μL forward/reverse primer and UV-treated H_2_O. Primer sequences can be found in Supplementary Table 1. Gene expression was analysed in duplicate on three separate occasions, no reverse transcription (NRT) and no template controls (NTC) were included throughout. Gene expression was normalised to the *ACT1* housekeeping gene and calculated using the ΔΔCt method.

### Strain and culture conditions for transcriptional analysis

*C. albicans* and *L. crispatus* type strains SC5314 and ATCC 33820, respectively, were used for transcriptional analysis in this study. For biofilm formation, overnight cultures of *C. albicans* and *L. crispatus* were grown in YPD and de Man, Rogosa and Sharpe (MRS, Merck UK) media, respectively, under appropriate culture conditions. Cultures were washed twice with PBS and standardised in 1:1 broth to 1×10^6^ CFU/mL for *C. albicans* and 1×10^7^ CFU/mL for *L. crispatus*.

### In vitro transcriptomic analysis of *C. albicans* interactions with *L. crispatus*

Overnight cultures were standardised as described above and *C. albicans* biofilms grown in T-75 cell culture flasks (Corning, USA) for 4 h in 5% CO_2_. Following incubation, media was removed, and biofilms washed before *L. crispatus* was added for an additional 2, 4 or 20 h. At each time point the media was removed and biofilms washed with PBS before being scraped in to 1 mL of RNA*later* (ThermoScientific, Loughborough, UK). Spent media from biofilms was retained and pH monitored throughout the experiment. RNA was extracted from microbial biofilms using the RiboPure™ RNA Purification Kit for yeast (ThermoScientific, Loughborough, UK), following manufacturer’s instructions. Integrity of RNA was assessed using a Bioanalyser system and genome-wide *Candida* transcripts sequenced using the Illumina NOVASeq6000 sequencing platform (Edinburgh Genomics). A summarised illustration of the experimental and bioinformatics pipelines can be found in Supplementary Figure 1.

### Investigating the probiotic potential of L. crispatus in a complex biofilm model

Mature 11-species biofilms were formed as described previously by our group with slight modifications [33]. Overnight broths of each organism were standardised in 1:1 medium prior to addition to the biofilm on hydroxyapatite (HA) discs. For probiotic treatment, each biofilm was treated bi-daily (12 h intervals) with 5×10^7^ CFU/mL of *L. crispatus* for 5 min before treatment was removed, biofilms washed 3 times in PBS and fresh media replaced. On day 3, after 4 probiotic treatments, 48 h biofilms were analysed for compositional analysis. On day 5, following 8 probiotic treatments, 96 h biofilms were removed for compositional analysis. At each timepoint, biofilms were washed 3 times and sonicated in 1 mL of PBS at 35 KHz for 10 min to remove biomass. Sonicates were split in two, with one sample having 5 μl of 10 mM propidium monoazide (PMA) added to it for the quantification of live *C. albicans* DNA. PMA is DNA-intercalating dye used to bind DNA from dead cells or those with a compromised membrane following exposure to a halogen light [34] [35]. This covalent bonding prevents downstream amplification of DNA in qPCR, resulting in amplification of DNA from live cells only. The other sample with no PMA allowed for quantification of total *C. albicans* DNA per biofilm. All samples were incubated in the dark for 10 min then placed on ice and exposed to a 650W halogen light for 5 min.

DNA was then extracted using the QIAmp DNA mini kit, as per manufacturer’s instructions (Qiagen, Crawley, UK). To a 20 μL qPCR reaction, 1 μL of biofilm DNA was added to: 10 μL Fast SYBR™ Green Master Mix, 1 μL of 10 μM *C. albicans* forward/reverse primers and UV-treated nuclease-free water. Primer sequences can be found in Supplementary Table 1. The following thermal profile was used: 95°C for 2 min, 40 cycles of 95°C for 3 s followed by 55°C for 30 s. Samples were assessed in duplicate from 2 separate experiments. Fungal CFE/mL were then calculated as described above.

### Statistical analysis

Microbiome figures were created using MicrobiomeAnalyst [36]. Transcriptome pipeline figures were constructed using BioRender and differential gene expression plots using the DESeq2 package in RStudio. Gene ontology networks were constructed using ClueGO software, available through Cytoscape [37]. All other figures and analyses were performed in GraphPad Prism (Version 8, La Jolla, California, USA).

## Results

Swab and lavage samples were collected from healthy individuals (n=60) and women suffering from RVVC (n=40). Demographic data collected through the patient questionnaire as well as pH and IL-8 levels are reported in Supplementary Table 1. Detection of increased cytokine levels in RVVC was confirmatory of clinical diagnosis (P < 0.0001).

The mechanisms behind the shift in *Candida* from commensal to pathogenic yeast seen in RVVC onset remains poorly understood. The cause of infection is likely multifactorial and here we aimed to assess whether fungal or bacterial load may influence disease. To this end, levels of *Candida* present in the clinical samples were measured by determining colony forming units (CFU/mL), in addition to total yeast and bacterial DNA quantified by qPCR (CFE/mL) (Figure 1). Numbers of CFU/mL showed a significant increase in *Candida* load in samples from patients with RVVC from around 2×10^3^ CFU/mL in healthy patients to 2.3×10^4^ CFU/mL in diseased (P<0.01) (Figure 1a). Similarly, when assessed molecularly, a significant increase of around 1.3-log was observed in patients with RVVC to 3.8×10^5^ CFE/mL (P<0.05), as shown in Figure 1b. This data suggests that increased levels of *Candida* could be a contributing factor to disease pathology and may be an indicator for VVC onset and subsequent recurrence. Additionally, *Candida* load was observed with respect to patient metadata; Although a slight reduction of between 0.5 and 1-log is seen in patients with disease for over 7 months, this was not significant (data not shown). An increase in fungal load from 1×10^4^ to 5×10^4^ CFU/mL was observed in women who had received treatment for RVVC longer than 1 month prior to sample collection (P<0.05) (Supplementary Figure 2). Bacterial load was found to be comparable at ~8×10^7^ CFE/mL between the two disease states (Figure 1c). Finally, quantities of bacterial to fungal DNA were found to be lower in RVVC, confirming that patients with disease have a higher fungal burden (Figure 1d).

**Figure 1:**
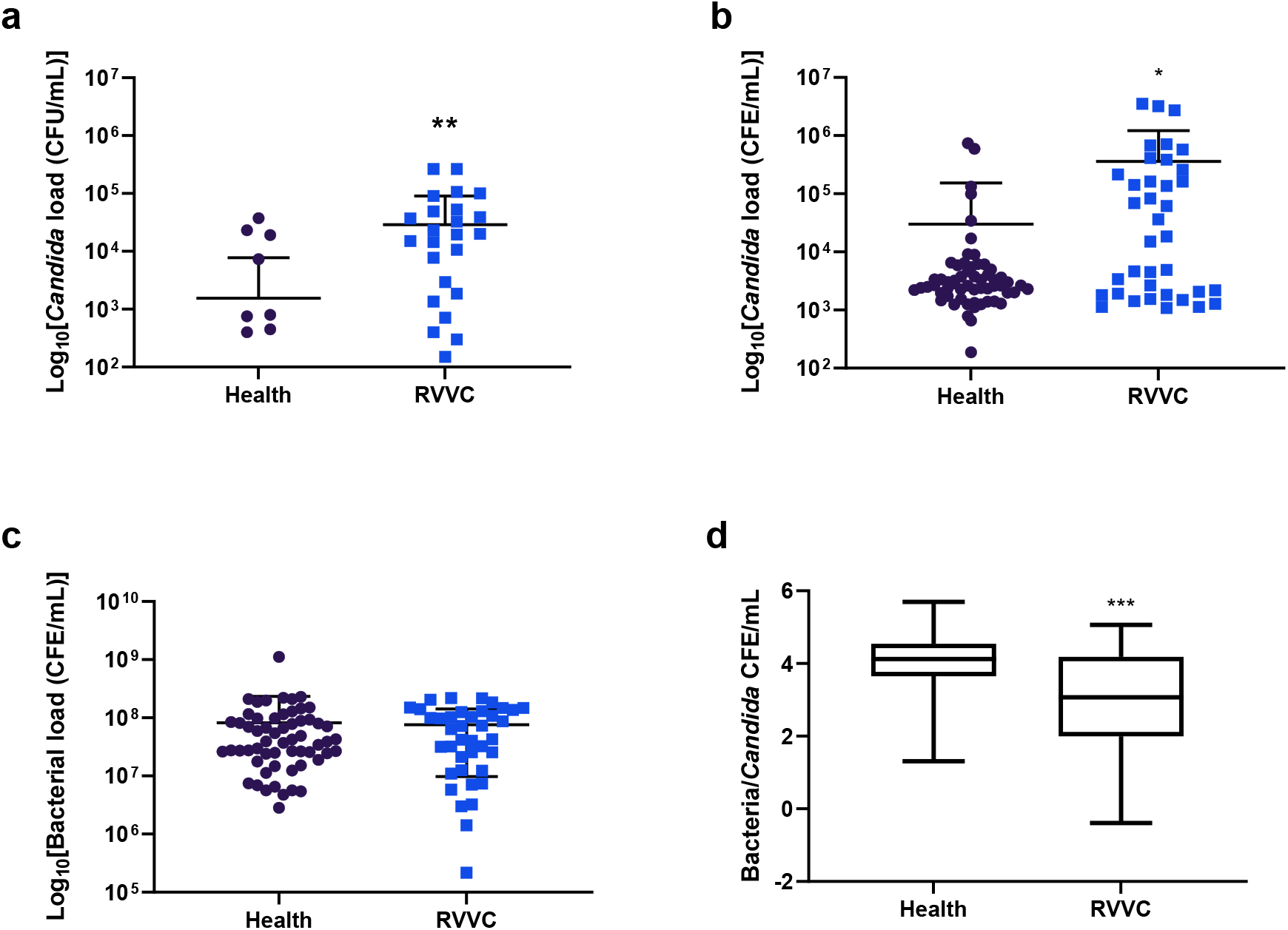
Fungal burden is elevated in women with RVVC while bacterial load remains unchanged. To asses fungal load, patient lavage was incubated on *Candida* Chromogenic agar plates and colonies counted after 48h **(a)**. For calculation of CFE/mL, the ITS region of *Candida* was amplified using genus-specific primers **(b)**. Levels of bacteria DNA were also assessed molecularly by amplification of the 16S rRNA region **(c)**; Correlation between bacterial and fungal burden was also observed **(d)**. Data represents the mean ± SD (*, P < .0.05, **, P < 0.01); Statistical significance was calculated using unpaired t-tests with Welch’s correction as data did not share equal standard deviations.

To observe bacterial taxa present in healthy women and those with RVVC, we performed 16S rRNA sequencing on DNA extracted from swab samples (Figure 2). At family-level, bacterial taxa present were found to be very similar between health and RVVC (Data not shown). At genus-level, a *Lactobacillus*-dominated environment accounting for up to 75% of the microbiome was observed in both cohorts with similar levels of diversity and vaginal anaerobes: *Gardnerella a*nd *Prevotella* (Figure 2a and 2b). *Atopobium* was found to be slightly higher in healthy patients and *Bifidobacterium* slightly lower. To ensure identity specificity at species-level, we analysed the microbiome of a subset of samples using the Oxford Nanopore Technologies MinION sequencer [38]. Although there is some fluctuation in abundances, this analysis confirmed accurate identification of *Lactobacillus* species by Illumina sequencing (Supplementary Figure 3). When viewed at species-level, subtle differences in bacterial taxa were observed between health and RVVC, particularly amongst *Lactobacillus* species (Figure 3). Most notably was the reduction in levels of specific *Lactobacillus* species including *L. jensenii* and, to a greater extent, *L. crispatus* which fell from 44% in health to 30% in RVVC (Figure 3a and 3b). Interestingly, this reduction was coupled with an increase in *L. iners* from just 19% in health to 40% in RVVC. Additionally, when predicted using random forest plots, levels of *L. iners* were suggested to be the most distinct between health and RVVC (Figure 3c).

**Figure 2:**
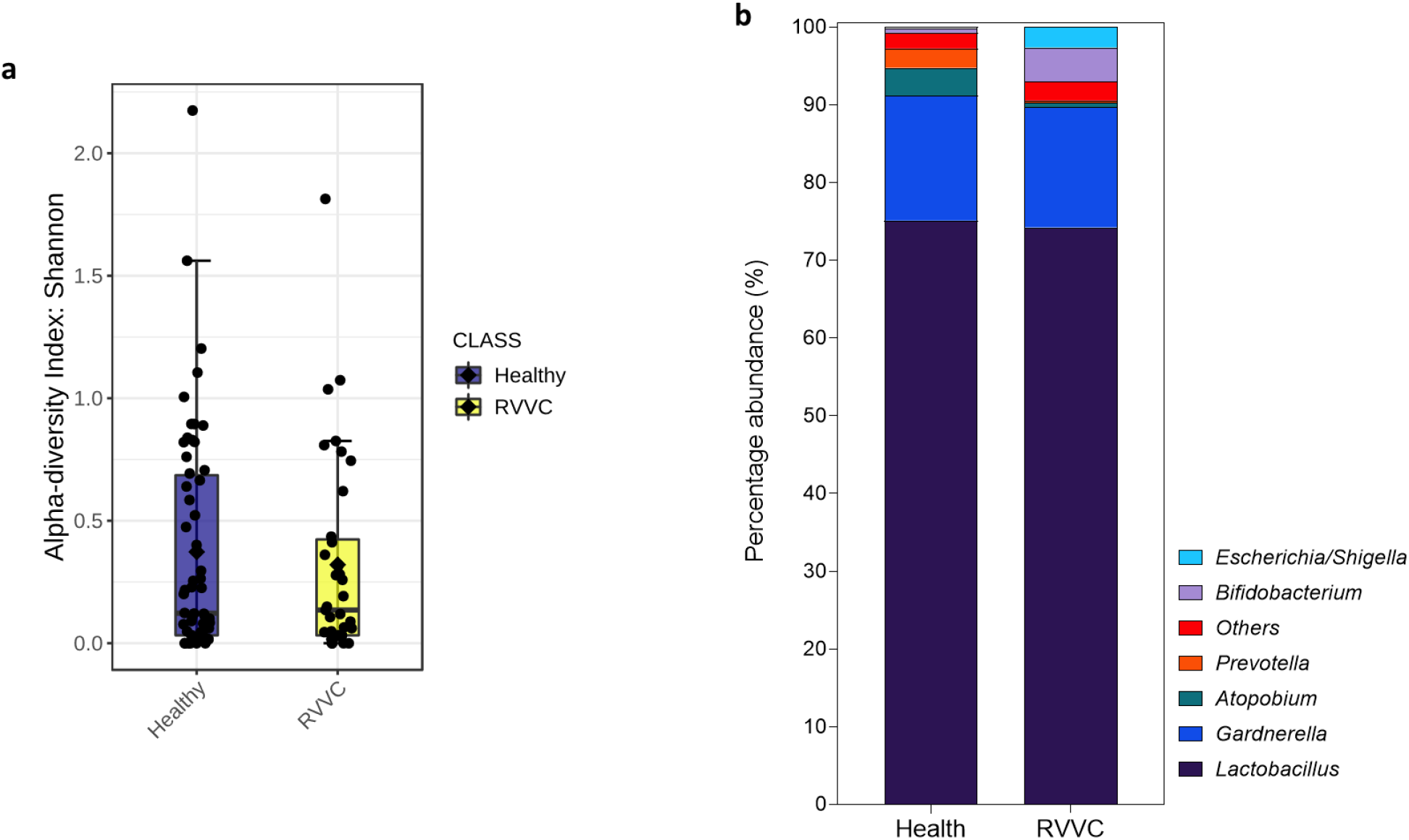
Bacterial genera present in health and RVVC. DNA extracted from swab samples was used for 16S rRNA Illumina sequencing (n=100). Bacterial diversity measured by Shannon index **(a)** and genus-level taxa identification and percentage abundance of microbial populations present **(b)** in health and RVVC.

**Figure 3:**
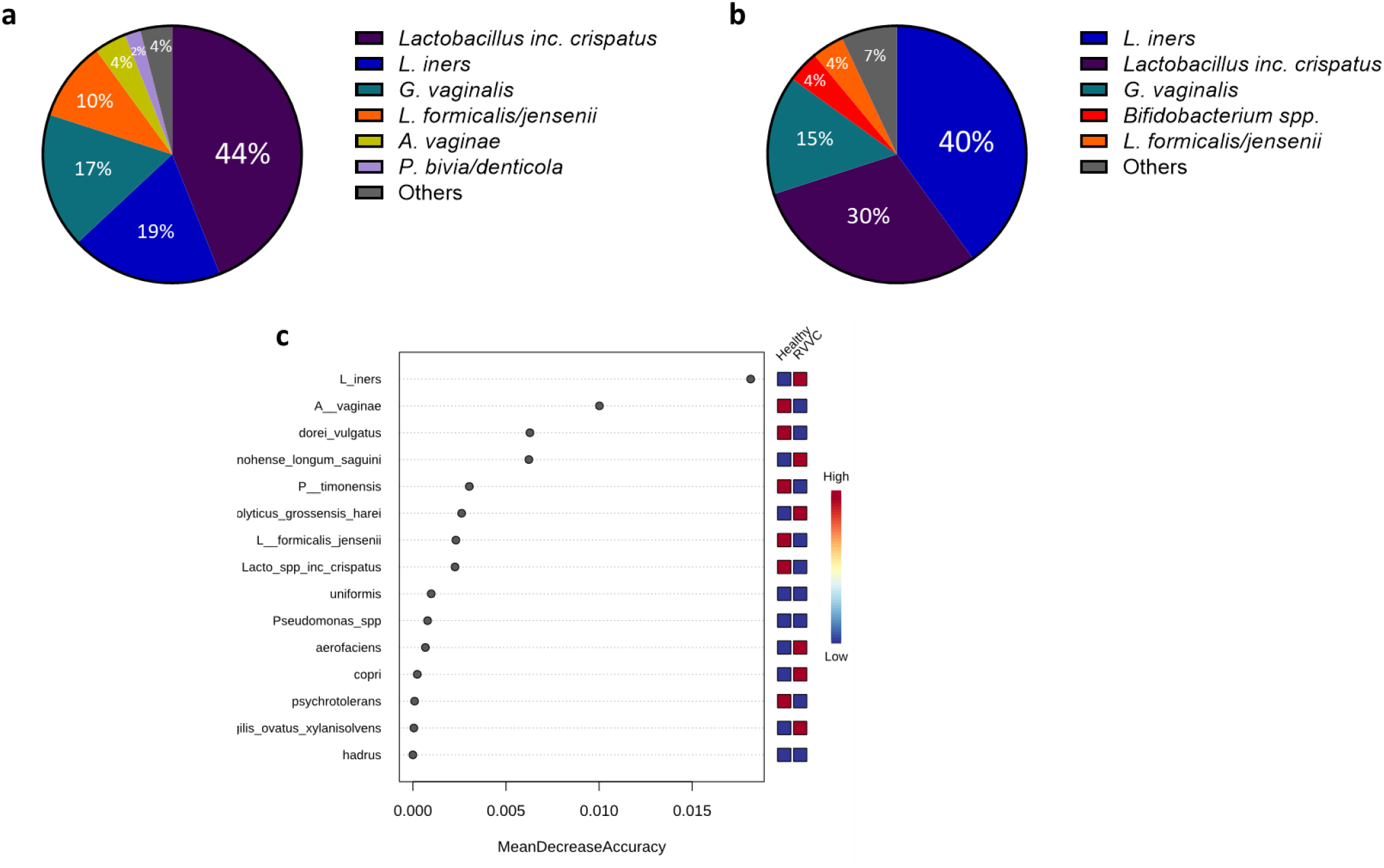
Hydrogen peroxide-producing lactobacilli strains are reduced during RVVC infection resulting in an *L. iners*-dominated microbiome. Species-level identification of bacterial taxa present in health **(a)** and RVVC **(b)**. Random forest plot showing the most distinct species-level taxa present between health and RVVC **(c)**

To investigate the influence of bacterial taxa further, microbial populations were observed with respect to patient metadata (Figure 4). In patients with culturable *Candida* compared with those who were culture-negative, a reduction in *Lactobacillus* species including *L. crispatus* from 44% to 29%, coupled with an increase in *L. iners* from 23% to 35% was observed, similar to the RVVC profile (Figure 4a). Further, *L. iners* was predicted to be the second-most likely organism to define presence or absence of *Candida* using random forest analysis to identify important features (Figure 4c). When observed with respect to the length of time patients had suffered from RVVC, an intermediate profile between health and disease was seen in patients with disease for less than 6 months, showing a slight reduction in *L. crispatus* from 45% to 40% (Figure 4c). Patients with disease for between 6-12 months, conversely, had a similar profile to that seen in RVVC with increased levels of *L. iners* from 19% to 44% (Figure 4d).

**Figure 4:**
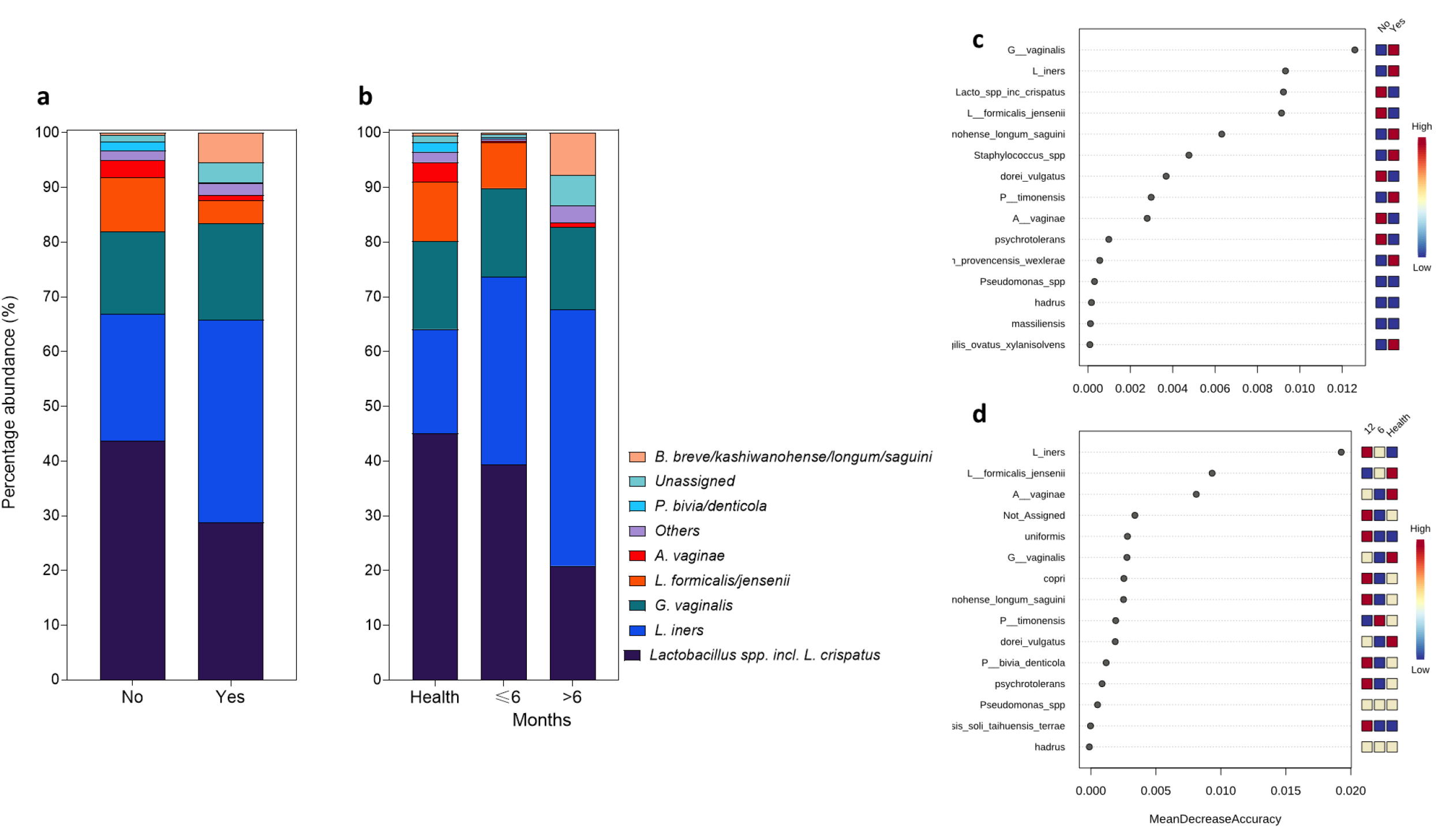
Species-level taxa abundance relative to patient metadata. Species-level bacterial taxa based on presence/absence of *Candida* **(a)** and length of time with disease **(b)**. Random forest plots showing distinct bacterial taxa present in each analysis **(c** and **d**).

*Candida* clinical isolates were obtained from lavage samples by culture on *Candida* chromogenic agar. Following incubation, a total of 33 isolates were obtained, 9 from healthy individuals and 24 from patients with RVVC. Isolate identification was confirmed using MALDI-TOF (Figure 5a). Consistent with previous RVVC studies, *C*. *albicans* was found to account for 73% of the *Candida* species isolated within our patient subset. Other NAC species, more commonly associated with RVVC, accounted for the remaining 27%. Following identification, biofilm forming capabilities of clinical isolates was assessed (Figure 5b). *C. albicans* was capable of forming dense biofilms with clear heterogeneity between isolates. Isolates were grouped as low biofilm formers (LBF) if the absorbance reading of their total biomass fell below the first quartile (< 0.185). Similarly, high biofilm formers (HBF) were observed when total biomass was above the third quartile (> 0.854). Intermediate biofilm formers (IBF) had biomass readings between these values. Little biomass was observed in NAC species. Additionally, when lavage from a RVVC patient with a *C. albicans* isolate capable of dense biofilm formation was viewed microscopically, clear hyphal formation and bacterial-yeast aggregates could be seen (Figure 5c). To further investigate the potential impact of *Candida* biofilms *in vivo*, expression levels of key biofilm-related genes were measured in patient lavage. This data shows detectable levels of expression of genes involved in hyphal morphogenesis, biofilm formation and pathogenesis in *Candida* present in patients with RVVC (Supplementary Table 3).

**Figure 5:**
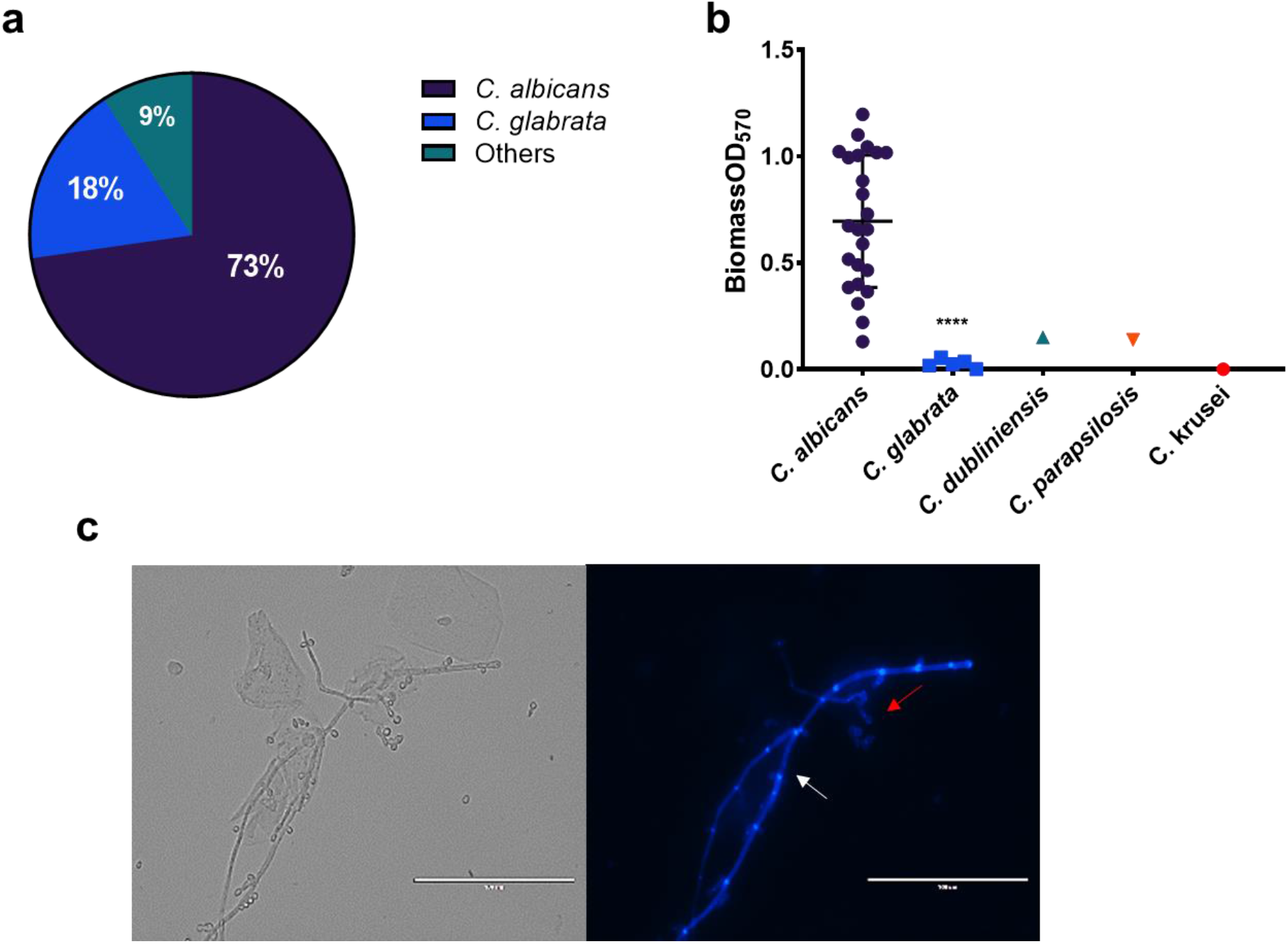
Vaginal *Candida* isolates from patients with RVVC are capable of heterogeneous biofilm formation. A total of 33 *Candida* clinical isolates were isolated from lavage samples. MALDI-TOF was used to identify each isolate and assess species distribution (Others are comprised of 1 isolate of: *C. dubliniensis, C. parapsilosis* and *C. krusei*) **(a)**. *Candida* isolates from health (n=9) and RVVC (n=24) were assessed for biofilm forming capabilities by crystal violet staining **(b)**. Vaginal lavage was stained with Calcofluor White (CFW) for 1 h at 37°C before imaging **(c)**. Images are representative of *C. albicans* aggregates and hyphae observed in lavage from a patient with a HBF isolate. White arrows represent pseudo-hyphal/hyphal formation and red arrows depict cell aggregates. Data represents the mean ± SD (****, P < 0.0001), statistical analysis was performed using unpaired t-tests.

To assess potential antagonism between *Lactobacillus* and *C. albicans*, 7 *Lactobacillus* species were selected and their effects on *C. albicans* biofilm formation in co-culture observed (Figure 6). When grown together for 24 h, a reduction in biomass was observed in all *Lactobacillus* species from an absorbance reading of 3.0 to around 2.5. This reduction was particularly prominent in *L. rhamnosus* where biomass was reduced to an absorbance value of 2.0 (Figure 6a). However, when *C. albicans* was allowed to form a biofilm before addition of *Lactobacillus*, this effect was less pronounced, and in some cases, absent (Figure 6b). To further analyse this antagonistic effect, *C. albicans* biofilm-related gene expression was assessed after co-culture with *L. rhamnosus* and *L. iners*. The two organisms interacted with *C. albicans* differently, with *L. rhamnosus* down-regulating all biofilm-related gene expression after incubation for 20 h and 24 h (Figure 6c). Gene expression was reduced by ~4-fold for HWP1 and between 0.5 to 1-fold for ALS3 and ECE1 in both growth conditions. *L. iners* down-regulated expression of *ECE1* and, to a lesser extent, *HWP1* after incubation for 24 h. However, when added to a pre-existing *C. albicans* biofilm, *L. iners* resulted in up-regulation of *ECE1* and *ALS3*.

**Figure 6:**
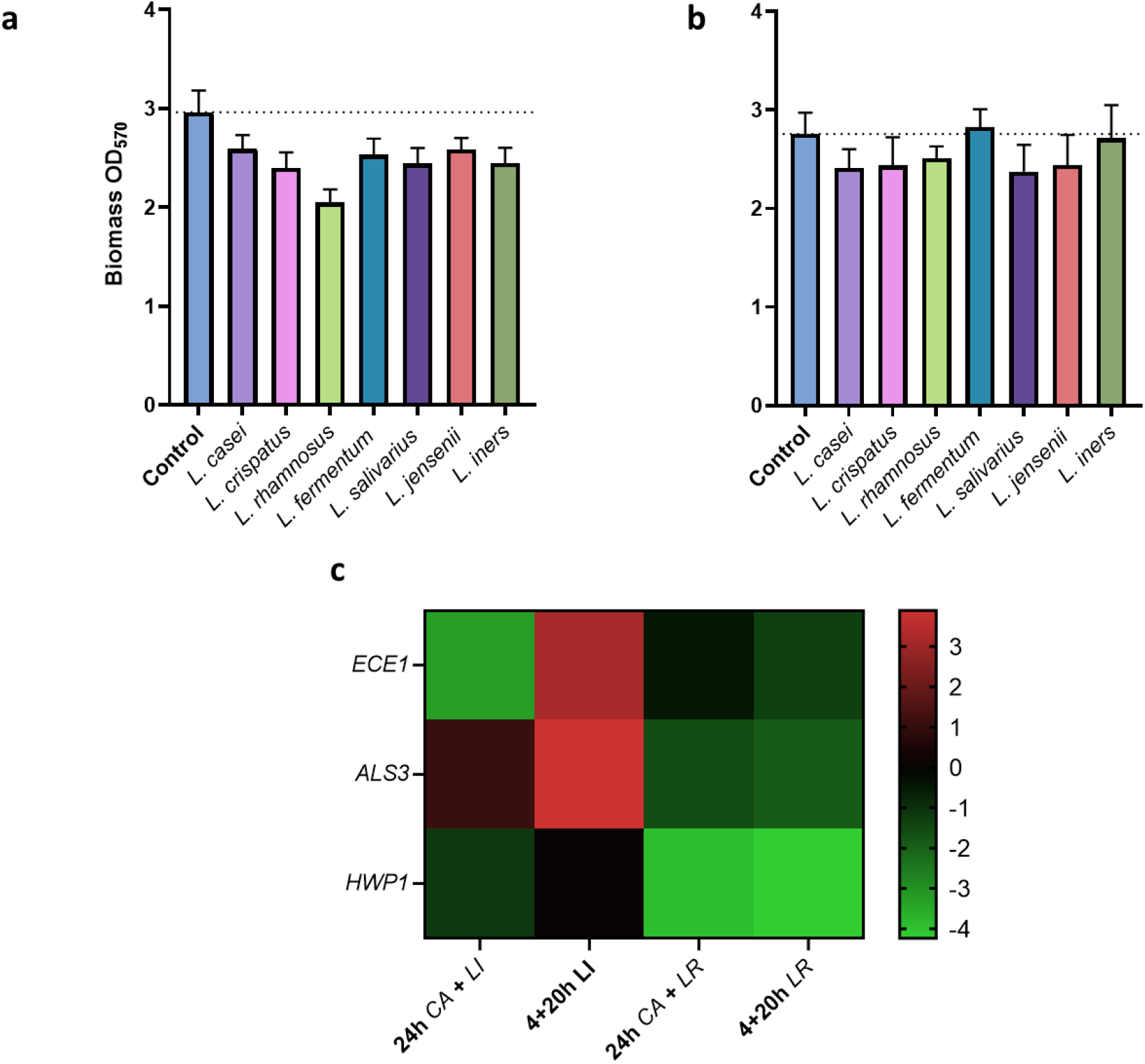
*Lactobacillus* species display antagonism with *C. albicans in vitro*. To observe inhibitory effects of *Lactobacillus* against *C. albicans* biofilm formation, *C. albicans* and a panel of *Lactobacillus* species were co-cultured together in THB/RPMI (1:1) media in 5% CO_2_ either for 24h **(a)** or *C. albicans* was grown for 4h prior to addition of *Lactobacillus* species for 20h **(b)**. *C. albicans* biofilm-associated gene expression was measured in the presence *L. rhamnosus,* which is associated with ‘health’, and *L. iners,* which is hypothesised to indicate dysbiosis. The mean log fold-change relative to single species *C. albicans* biofilms is shown **(c)**. Data represents the mean + SD.

In this study, we aimed to utilise whole-genome transcriptional sequencing of *C. albicans* following co-culture with *L. crispatus* to elucidate intimate interactions between the two organisms (Supplementary Figure 1). *L. crispatus* was selected for RNA sequencing analysis as *L. rhamnosus* is not a commensal of the vaginal environment and is only capable of transient colonisation. Further, *L. crispatus* is one of the most prevalent *Lactobacillus* species in the vaginal microbiome and its antagonism with *C. albicans* has been displayed in previous studies [39]. Initially, multivariate analysis by Principal Component Analysis (PCA) was unable to identify distinct clusters of gene expression in early dual-species biofilms (Supplementary Figure 5). Subsequently, there was no significant up or down-regulation in early dual species biofilm transcripts compared to single species controls (P>0.05). For this reason, further analysis compared expression in mature 24 h single and dual species biofilms only. Initially, differential expression analysis was performed in R using the DESeq2 package to observe transcriptional changes between different biofilms and time-points (Figure 7). Heatmap analysis of normalised Log_2_ fold change in gene expression between single and dual species 24 h biofilms with *L. crispatus* revealed up-regulation of many genes involved in amino acid biosynthesis and breakdown in dual-species biofilms (*ARG8, ILV1, HIS5*) (Figure 7). Interestingly, the *PRY1* gene which codes for a secreted protein associated with virulence in the presence of lactate in *C. albicans* was found to be downregulated in dual species biofilms (Log_2_ fold change = −5.35). A list of some of the key genes and their function can be found in Table 1. Gene ontology (GO) term analysis was utilised to determine the functionality of differentially expressed genes in single and dual species 24 h biofilms. Differentially expressed genes were assigned to GO terms based on function. Genes expressed in *L. crispatus* dual-species biofilms were mainly responsible for amino acid biosynthesis and some metabolic and transaminase activity (Figure 8a and 8b). Despite upregulation of these amino acid biosynthesis and metabolism pathways in *C. albicans*, expression of *BAT21* and *ILV1* in dual-species biofilms suggests *C. albicans* was in a state of amino acid starvation (Figure 8c).

**Table 1:**
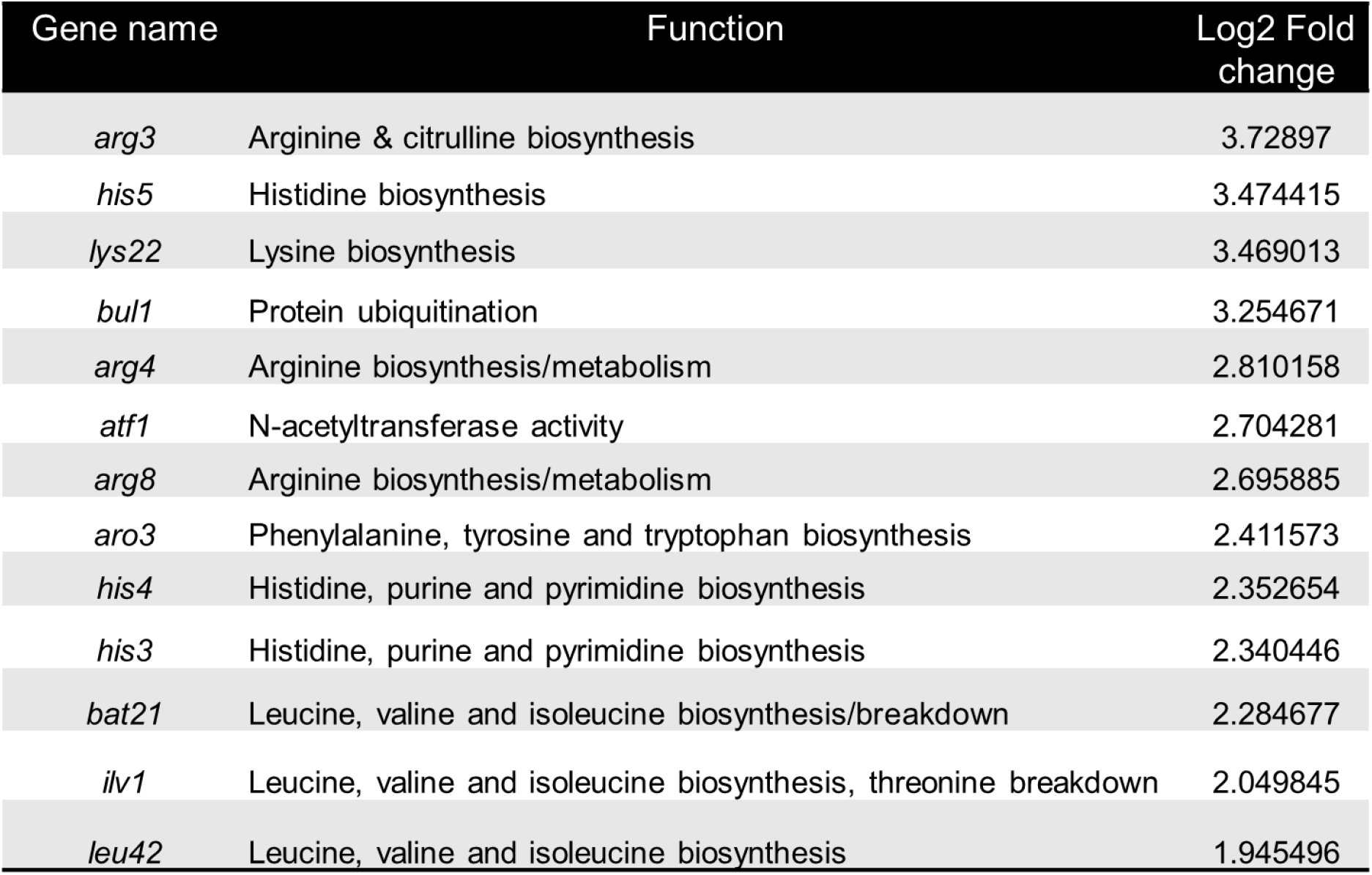
Up-regulated genes in 24h dual-species biofilm associated with amino acid biosynthesis and/or breakdown.

| Gene name | Function | Log2 Fold change |
| --- | --- | --- |
| *arg3* | Arginine & citrulline biosynthesis | 3.72897 |
| *his5* | Histidine biosynthesis | 3.474415 |
| *lys22* | Lysine biosynthesis | 3.469013 |
| *bul1* | Protein ubiquitination | 3.254671 |
| *arg4* | Arginine biosynthesis/metabolism | 2.810158 |
| *atf1* | N-acetyltransferase activity | 2.704281 |
| *arg8* | Arginine biosynthesis/metabolism | 2.695885 |
| *aro3* | Phenylalanine, tyrosine and tryptophan biosynthesis | 2.411573 |
| *his4* | Histidine, purine and pyrimidine biosynthesis | 2.352654 |
| *his3* | Histidine, purine and pyrimidine biosynthesis | 2.340446 |
| *bat21* | Leucine, valine and isoleucine biosynthesis/breakdown | 2.284677 |
| *ilv1* | Leucine, valine and isoleucine biosynthesis, threonine breakdown | 2.049845 |
| *leu42* | Leucine, valine and isoleucine biosynthesis | 1.945496 |

**Figure 7:**
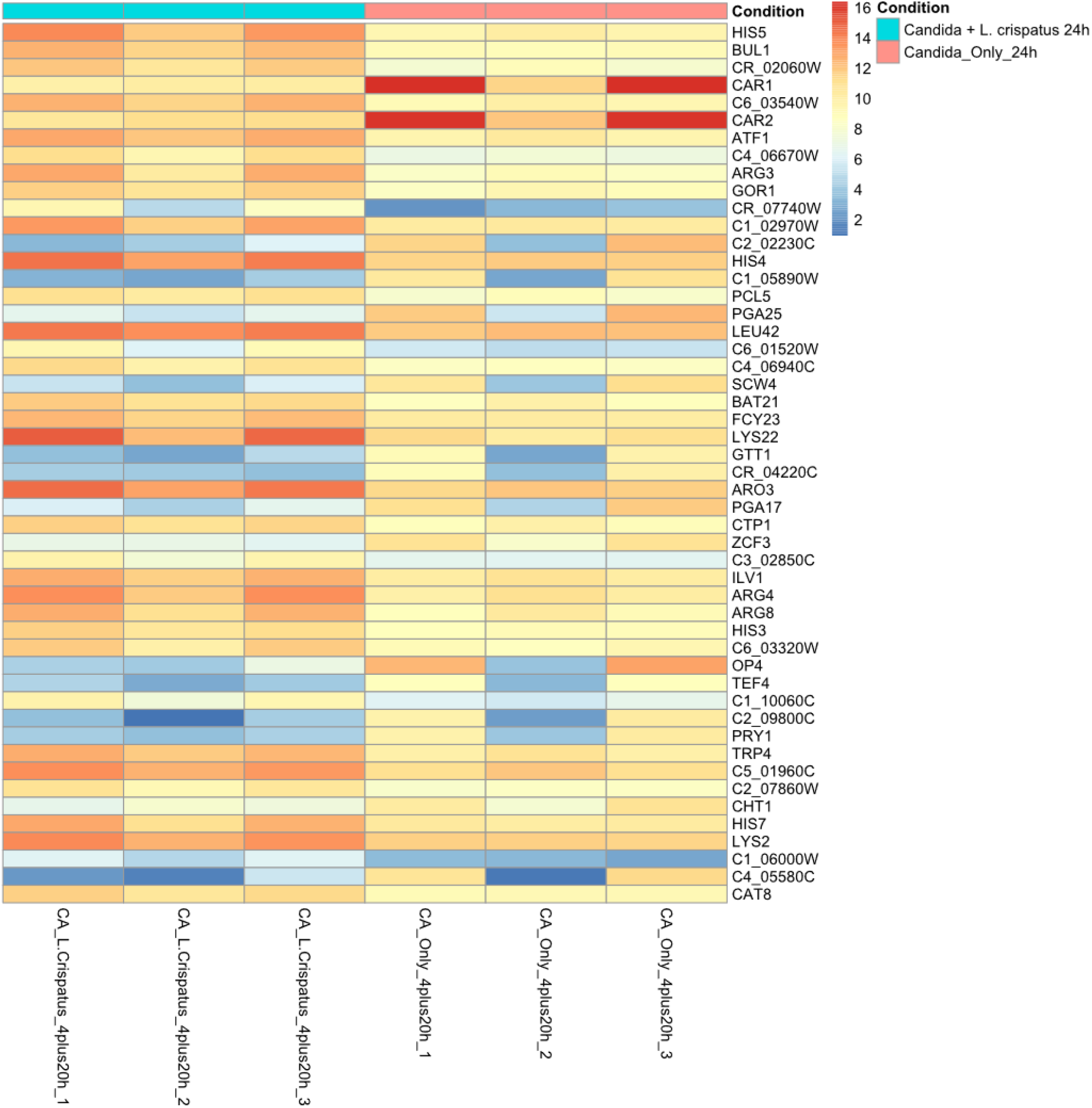
Differential expression analysis of *C. albicans* single and *C. albicans + L. crispatus* dual species biofilms. Heatmap displaying the top 50 significantly differentially expressed genes in *C. albicans* between single and dual species 24h biofilms (P < 0.05).

**Figure 8:**
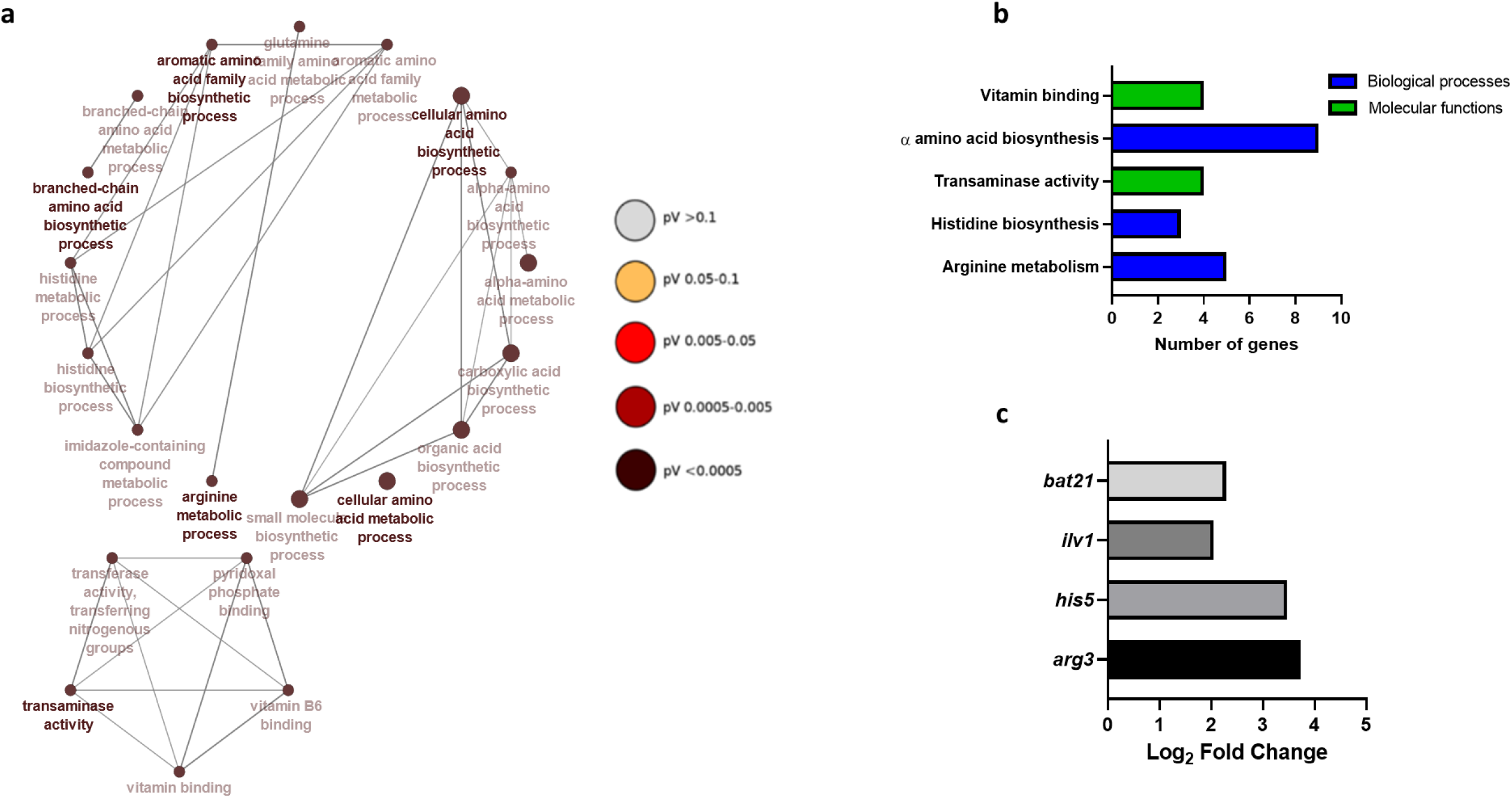
Gene networks of Gene Ontology (GO) terms and up-regulated genes in dual species biofilms. Constructed gene networks of GO terms in 24h biofilms with *L. crispatus* **(a)**. Important biological, cellular and molecular functions in dual species biofilms with *L. crispatus* **(b)**. Log_2_ fold change of key gene expression in *C. albicans* from single to dual species biofilms with *L. crispatus* **(c)**. Nodes are coloured by significance. All GO terms have an adjusted P value < 0.05. Networks were created using ClueGO.

Given the antagonism observed between the two organisms, we then aimed to investigate the *in vitro* probiotic potential of *L. crispatus* against *C. albicans* infection in a complex biofilm model (Figure 9). After 2 consecutive days of probiotic treatment, a slight reduction in total and live *C. albicans* composition within the biofilm is seen, however this was not significant (P = 0.55, P = 0.16, respectively) (Figure 9a and 9b). Following a 4-day treatment regimen with *L. crispatus*, total *C. albicans* composition decreased (P < 0.05) with reduced levels of live fungal DNA. When assessing the total fold reduction in *C. albicans* from untreated biofilms, the greatest probiotic effect is seen at 48h post-treatment (Figure 9c).

**Figure 9:**
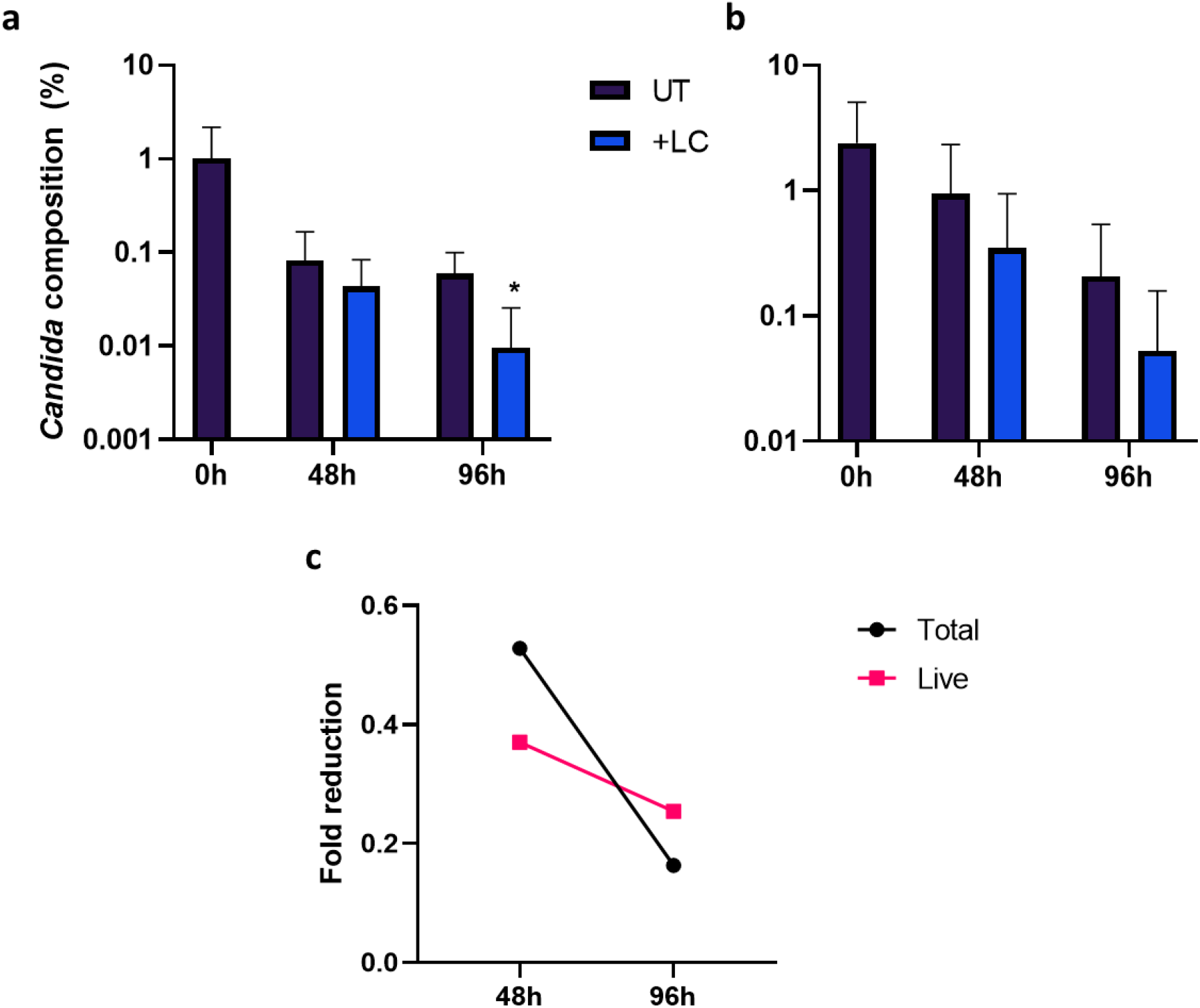
Bi-daily addition of *L. crispatus* reduces *C. albicans* load within a complex biofilm model *in vitro*. The potential probiotic properties of *L. crispatus* against *C. albicans* were assessed in a 11-species biofilm model treated twice daily with *L. crispatus.* Live/Dead qPCR allowed for quantification of percentage composition of total **(a)** and live **(b)** *C. albicans* DNA within the biofilm. Average fold change in the *C. albicans* percentage composition from the untreated 11-species biofilm is also shown **(c)**. Data represents mean + SD. Statistical analysis was performed using paired t-tests comparing raw CFE values (*, P < 0.05).

## Discussion

Unlike systemic and oral candidiasis, RVVC affects immunocompetent women with an incidence rate of up to 8%, hence it is the most prevalent human infection caused by the pathogenic yeast, with approximately 140 million cases annually [40]. Despite this, the disease remains largely understudied in the field of women’s health. A better understanding of RVVC at a molecular level may be crucial in unearthing potential targets for much-desired therapeutics. This study aimed to asses a panel of 100 clinical isolates to investigate fungal influence, changes in bacterial communities and to use an RNA sequencing approach to analyse potential interkingdom interactions contributing to disease pathology.

The estimated vaginal commensal carriage rate of *Candida* is 33% [41]. We observed the presence of *Candida* both by culture and by qPCR and found 15% and 60% of healthy and RVVC samples had culturable *Candida*, respectively, and all but 2 samples were positive by qPCR (Supplementary Table 1). An increased pH in RVVC, although not diagnostic, is seen in practice, likely due to a loss of *Lactobacillus* species [42]. This was reflected in our study where vaginal lavage measured at a pH of approximately 5, higher than the reported 4.5 of health. As expected, the inflammatory biomarker, IL-8, and levels of *Candida* DNA were found to be significantly higher in patients with RVVC, confirmatory of clinical diagnosis (Figure 1 and Supplementary Table 1). It is important to note that these clinical samples are from women regularly attending sexual health clinics, and those who have suffered from RVVC for longer may be more likely to be receiving antifungal treatment. This should be taken in to account when interpreting *Candida* CFE/mL data observed in this study as untreated RVVC may present differently.

Unlike vaginal infections such as BV, the microbial communities present during VVC have been shown to be similar with those present in health at the phylum and genus-level [43, 44]. Our study confirms these findings where we report a *Lactobacillus*-dominated population with vaginal anaerobes including *Gardnerella*, *Prevotella* and *Atopobium*, with no significant differences in diversity or composition between the two cohorts at the genus-level (Figure 2). This suggests that the functional capacity of the bacterial species found in health and RVVC may play a more crucial role in disease pathology. A limitation of our microbiome analysis was the inability to distinguish between some *Lactobacillus* species (including *L. acidophilus, L. casei* and *L. gasseri*). For this reason, these species are grouped as ‘*L. spp*, incl. *L. crispatus’*. However, sequencing of a subset of samples using the MinION sequencer confirmed accurate identification of *Lactobacillus* to species-level (Supplementary Figure 3). We observed a reduction in specific *Lactobacillus* species, including those associated with maintaining health due to their ability to produce L-lactic acid and H_2_O_2_, such as *L. crispatus* and L*. jensenii* (Fig 4). This reduction coupled with an increase of *L. iners* has been shown previously and is thought to be indicative of vaginal dysbiosis [43]. We investigated this phenomenon further with respect to individual patient metadata (Fig 5). We report that this loss of health-associated *Lactobacillus* species and increase of *L. iners* is also seen specifically in patients with positive *Candida* cultures and those who have suffered from RVVC for >6 months. These findings may be important for future studies investigating RVVC. It is hypothesised that changes within *Candida* allow for it to switch from asymptomatic commensal to pathogenic yeast. It may now be more important to study RVVC with specific focus on the microbes present during RVVC, specifically *Lactobacillus*, and interkingdom interactions which may influence this pathogenic switch in *Candida*.

Consistent with epidemiological reports, we found >70% of isolates from our cohort were *C. albicans* with the remainder comprised primarily of *C. glabrata* (Figure 6). Though *C. albicans* has been shown to form biofilms on vaginal mucosa and *Candida* clinical isolates shown to form biofilms *in vitro*, the presence of biofilms during RVVC infection is still disputed [12, 13, 45]. Despite evidence to the contrary, our study has shown heterogeneous biofilm formation in *C. albicans* RVVC isolates and shows clear visualisation of *C. albicans* hyphae and aggregates in lavage fluid from a patient with RVVC (Figure 6). Therefore, *Candida* biofilm formation within the vagina should not be discounted as a potential cause of failed clinical treatment and subsequent recurrence of disease. We also show the capacity of various *Lactobacillus* species to inhibit *C. albicans* biofilm formation when co-cultured for 24h (Figure 7). *L. rhamnosus* has been studied extensively for its potential as a probiotic and has been shown to prevent adhesion of *C. albicans* to mucosal surfaces [46]. Here we show the ability of *L. rhamnosus* to down-regulate *C. albicans* biofilm-related gene expression. In contrast, *L. iners* results in up-regulation of both *als3* and *ece1*. This is consistent with previous findings suggesting *L. iners* may be indicative of a dysbiotic vaginal environment and should not be considered as a probiotic intervention for C. albicans infections. and as such, should not be considered as a probiotic treatment for *C. albicans* infections [43].

*L. crispatus* is increasingly becoming recognised as a health-associated *Lactobacillus* species in the vaginal microbiome and may be important in preventing RVVC. In our transcriptomic experiment we aimed to elucidate possible antagonism between *C. albicans* and *L. crispatus* (Figure 7). Initially, differential gene expression analysis by Principal Component Analysis (PCA) was unable to identify distinct clusters of gene expression between early single and dual-species biofilms (Supplementary Figure 5). For this reason, subsequent analysis was performed comparing single and dual-species biofilms at 24h only. It has previously been shown in *Lactobacillus* clinical isolates that secreted metabolites can remain at low levels until up to 72 or 120h incubation, which may account for the delay in interaction seen here [47]. The *in vivo* vaginal pH is thought to be <4.5, however, in this dual species model the pH was sustained between 6-7 (Supplementary Figure 4). It is possible that in the presence of other lactic acid bacteria, the low environmental pH could affect *C. albicans* gene expression. Nonetheless, the stability of the neutral pH throughout this experiment dictates that any transcriptomic changes observed are not due to acidification of the media and are a result of interactions between *C. albicans* and *L. crispatus*.

Gene transcripts associated with amino acid biosynthesis and transaminase activity were prominent in *L. crispatus* dual-species biofilms (Figure 8). Transaminase activity is primarily associated with α-amino acid breakdown and biosynthesis. It has been shown in a model of *Saccharomyces cerevisiae, L. lactis* and *L. plantarum*, that in Nitrogen rich environments yeasts secrete an array of metabolites, primarily amino acids, thereby facilitating the growth of lactobacilli [48]. The upregulation of various amino acid biosynthesis processes in our study may be attributed to this nutrient cross-feeding. This may be a deliberate process by which *L. crispatus* is able to drive synthesis of amino acids in *C. albicans*, as well as suppressing *car1* and *car2*, associated with arginine breakdown, in order to sequester them for metabolism and facilitate its own growth. The upregulation of *bat21* and *ilv1* in dual species biofilms is associated with amino acid starvation in *C. albicans*, suggesting that despite the upregulation of various amino acid biosynthesis processes, *C. albicans* is unable to utilise them. This may be a contributing factor in the process by which *L. crispatus* is able to out-compete *Candida* during VVC infection and potentially re-establish a healthy microbiome. Furthermore, the *pry1* gene, a secreted glycoprotein associated with virulence and sterol binding in the yeast cell wall in the presence of lactate, was found to be downregulated in dual species biofilms [49]. Thus, *L. crispatus* may possess a mechanism by which it can suppress lactate-associated virulence in *C. albicans*. Interestingly, genes associated with resistance to lactic acid such as *mig1* and *cyb2* were not found to be differentially expressed in our co-culture study [50, 51]. This suggests a classical weak organic acid stress response in *C. albicans* is not being triggered by the presence of *L. crispatus* alone. Given that the media in this experiment was not acidified, this concludes that antagonism between *C. albicans* and *L. crispatus* is not purely due to a reduction in environmental pH. The influence of other secreted metabolites of *L. crispatus* such as H_2_O_2_ and bacteriocin-like substances on *C. albicans* requires further research to fully illustrate antagonism between the two organisms. These data show a potentially probiotic effect of *L. crispatus* against *C. albicans* in non-acidic environments. This antagonism is governed by the over-production of amino acids from *C. albicans* which may facilitate restoration of a healthy microbiome through lactobacilli proliferation.

Finally, we investigated the potential probiotic effect of *L. crispatus* against *C. albicans* within a complex biofilm model (Figure 9). Although studies have assessed the inhibitory effect of *L. crispatus* against *C. albicans* mono-species biofilms [39, 52, 53], to our knowledge, this is the first report of *L. crispatus* reducing *C. albicans* composition within a polymicrobial biofilm model *in vitro*. Our data supports other studies confirming the probiotic potential of *L. crispatus* against *C. albicans* biofilm infections.

In conclusion, our study suggests it may be important to view RVVC as a result of fluctuation in antagonistic interkingdom interactions between *Candida* and *Lactobacillus,* potentially as a result of *Candida* biofilm formation. Additionally, we show for the first time the potential of *L. crispatus* to re-establish a healthy vaginal environment through altered gene expression in *C. albicans* and the ability to reduce fungal composition in a complex biofilm model.

## Author Contributions

EM, LS, RK, CD and SW participated in study design and experimental procedures. EM, LS, RK and CD were responsible for reparation of the manuscript. RM and RT were responsible for clinical sample collection and collection of patient metadata. CW participated in study design and contributed to the manuscript. GR conceived the study, participated in study design and was responsible for producing the final manuscript. All authors have read and approved the final manuscript.

## Competing interests’ statement

The authors declare no conflict of interest.

## Notes

### Competing Interest Statement

The authors have declared no competing interest.

